# Inflammatory endothelial cells promote infiltration of antigen-licensed cytotoxic T cells in malignant gliomas after irradiation

**DOI:** 10.64898/2026.07.08.737259

**Authors:** Clara Tejido, Niklas Grassl, Nirmeen Elmadany, Jakob Rosenbauer, Zaira Seferbekova, Khwab Sanghvi, Dennis A. Agardy, Ralph Sinn, Theresa Bunse, Anna Mathioudaki, Kristine Jähne, Jana K. Sonner, Michael O. Breckwoldt, Abigail Suwala, Moritz Gerstung, Felix Sahm, Lukas Bunse, Michael Platten, Katharina Sahm

## Abstract

Insufficient T cell infiltration into malignant gliomas fundamentally limits the efficacy of adoptive T cell therapy. Here we show that fractionated irradiation overcomes this barrier by reprogramming the tumor endothelium towards an immune-recruiting interface.

Using complementary murine glioma models combined with adoptive T cell transfer, antigen-specific vaccination, and single-cell transcriptomic and T cell receptor profiling, we demonstrate that irradiation enhances the accumulation, clonal expansion, and effector differentiation of tumor-specific CD8^+^ T cells. Irradiated tumors showed increased T cell receptor clonality and local enrichment of proliferative effector CD8⁺ T cells with enhanced cytotoxic, interferon-responsive, and oxidative metabolic programs. Mechanistically, irradiation triggers a conserved interferon-driven endothelial program marked by antigen presentation and upregulation of adhesion molecules, including ICAM-1 and VCAM-1. This radiation-induced endothelial activation program preferentially seen in inflammatory endothelial subsets was conserved in human glioblastoma and linked to T cell recruitment and maintenance of activated CD8⁺ T cell states. Functionally, irradiation synergized with adoptive T cell transfer and antigen-specific vaccination to promote glioma-specific T cell accumulation and effector differentiation, improving tumor control and survival.

Together, these findings identify radiation-induced endothelial activation as a key regulator of T cell trafficking across the brain tumor vasculature highlighting the vascular niche as a critical determinant of immunotherapy efficacy and a rational target for combination strategies in glioblastoma.

## Introduction

Glioblastoma remains one of the most aggressive and treatment-refractory solid tumors, characterized by profound immunosuppression and limited responsiveness to immunotherapy (Bunse et al., 2025; Lim et al., 2018; Ostrom et al., 2020; Reardon et al., 2020). Despite the presence of tumor-infiltrating lymphocytes, effective anti-tumor immune surveillance is constrained by multiple barriers, including a hostile tumor microenvironment (TME), dysfunctional vasculature, and restricted immune cell trafficking across the blood–brain barrier (BBB) (Jung et al., 2021; Pichol-Thievend et al., 2024). In the central nervous system (CNS), immune cell entry is further regulated by specialized vascular and perivascular niches, including the endothelial layer and associated perivascular spaces, which actively control T cell adhesion, diapedesis, and positioning within the brain parenchyma (Engelhardt & Ransohoff, 2012; Engelhardt et al., 2017).

Radiotherapy is a central component of glioblastoma standard-of-care treatment and has traditionally been viewed as a purely cytotoxic modality; however, accumulating evidence indicates that ionizing radiation exerts broad immunomodulatory effects that extend well beyond direct tumor cell killing.

Radiation reshapes the tumor microenvironment through immunogenic cell death, enhanced antigen release and presentation, activation of innate immune sensing pathways, and modulation of stromal and vascular compartments. It triggers type I interferon responses, promoting dendritic cell activation, and fostering T cell priming and recruitment (Burnette et al., 2011; De Martino et al., 2021; Vanpouille-Box et al., 2017). At the same time, radiation can also induce counter-regulatory mechanisms, including immune checkpoint upregulation and recruitment of immune-suppressive cell populations, highlighting the context-dependent nature of its immunological consequences (van Hooren et al., 2023). Defining how these effects are coordinated across distinct cellular compartments remains critical for the rational design of combination therapies.

Recent work has emphasized the importance of resident and infiltrating T cell populations in mediating durable tumor control in glioma (Kilian et al., 2022; Luning et al., 2025; Watowich et al., 2023). Consistent with this, tumor-experienced T cells can acquire radiation-resistant and functionally adapted states that persist within irradiated tumors and contribute to local immune control (Belcaid et al., 2014; Dovedi et al., 2014; Zeng et al., 2013). However, the mechanisms by which irradiation promotes effective T cell accumulation and activation in the brain remain incompletely understood. In particular, how radiation influences the multistep process of T cell trafficking across the brain vasculature - including endothelial activation, perivascular retention, and re-entry into the parenchyma - remains poorly defined.

Beyond immune cells themselves, the tumor endothelium has emerged as a critical regulator of anti-tumor immunity in non-CNS tumors, in particular high endothelial venules (HEV) (Asrir et al., 2022; Peske et al., 2015). In the brain, endothelial cells form the blood-brain and blood-tumor barrier that impose additional constraints on immune cell entry and anti-tumor immune responses (Arvanitis et al., 2020). Unlike peripheral tissues, leukocyte extravasation in the brain occurs through a highly specialized, multi-step process involving endothelial cells, basement membranes, and perivascular spaces that act as transient niches guiding immune cell positioning before entry into the parenchyma(Engelhardt et al., 2017). Nevertheless, induction of HEV through combined antiangiogenic and anti-PD-L1 therapy has been shown to enhance antitumor immunity in glioblastoma similarly to non-CNS tumors (Allen et al., 2017).

While immunotherapeutic strategies such as tumor vaccination and adoptive T cell transfer are increasingly explored for the treatment of high-grade glioma (Brown et al., 2016; Bunse et al., 2026; Chih et al., 2025; Grassl et al., 2023; O’Rourke et al., 2017; Platten et al., 2021), the optimal scheduling and integration of radiotherapy to achieve synergistic anti-tumor immunity remain unclear. Although combining radiotherapy with antigen-directed immunotherapies has shown promise in preclinical models (Kim et al., 2017), the cellular and molecular mechanisms underlying these synergistic effects — particularly within the tumor vascular niche — are not fully defined.

In this study, we sought to dissect how irradiation reshapes the glioma immune microenvironment, with a particular focus on the coordinated responses of T cells and tumor endothelial cells. Using single-cell transcriptomic and T cell receptor profiling across multiple experimental glioma models, we investigated how radiation alone and in combination with antigen-specific immunotherapies modulates immune cell composition, functional states, and cell–cell interactions within the brain tumor niche. By specifically interrogating vascular–immune crosstalk at the BBB and perivascular interface, our work aims to define conserved radiation-induced programs that govern immune recruitment and activation in glioblastoma, thereby providing mechanistic insight to inform rational combination strategies for immunotherapy in the irradiated brain.

## Results

### Irradiation enhances tumor-specific CD8^+^ T cell accumulation and effector differentiation in malignant glioma

To study whether fractionated irradiation enhances antigen-specific T cell infiltration, we employed a GL261 glioma model expressing the tumor-associated antigen glycoprotein-100 (gp100) together with adoptive transfer of mCherry- and luciferase-expressing Pmel-1 CD8⁺ T cells, which express a transgenic T cell receptor specific for the gp100 epitope. After tumor establishment, mice received *ex vivo* primed gp100-specific CD8⁺ T cells intravenously. Fractionated irradiation of the tumor-bearing brain hemisphere in combination with adoptive T cell transfer (ACT) followed by *in vivo* T cell activation applying a gp100-targeting peptide vaccine extended median survival to >49 days compared to 21 days for ACT alone (Log-rank test p=0.00057) (Fig. 1A, Supplementary Fig. 1A-C). Bioluminescence imaging revealed 1.7-fold increase of T cell signals in the brain (7.53x10^5 vs 4.34x10^5 photons/s, p=0.006) (Fig. 1B-D) while draining lymph node and spleen signals remained unchanged (Fig. 1E-G, Supplementary Fig. 1D), suggesting enhanced recruitment and local accumulation of adoptively transferred T cells within irradiated tumors. In line, transferred gp100-reactive T cells isolated from irradiated tumors exhibited a proliferating effector phenotype suggesting robust *in situ* expansion: 22.1% Ki-67^+^, 10.6% IFNg^+^, 24.3% PD-1^+^, 4.2% TIM-3^+^, 17.4% LAG-3^+^ (all p<0.05 vs. controls) (Fig. 1F-M). Whole-brain tissue clearing and 3D lightsheet microscopy imaging revealed antigen-specific recruitment, with mCherry-expressing gp100-reactive CD8⁺ T cells localizing to gp100-expressing GL261 tumor cells in close proximity to the tumor vasculature, while T cells specific for the irrelevant antigen ovalbumin did not enter the tumor core (Figure 1 N, O). Furthermore, the proportion of gp100-reactive adoptively transferred T cells to endogenous T cells was higher in irradiated tumors compared to non-irradiated tumors (Supplementary Fig. 1E).

**Fig. 1:**
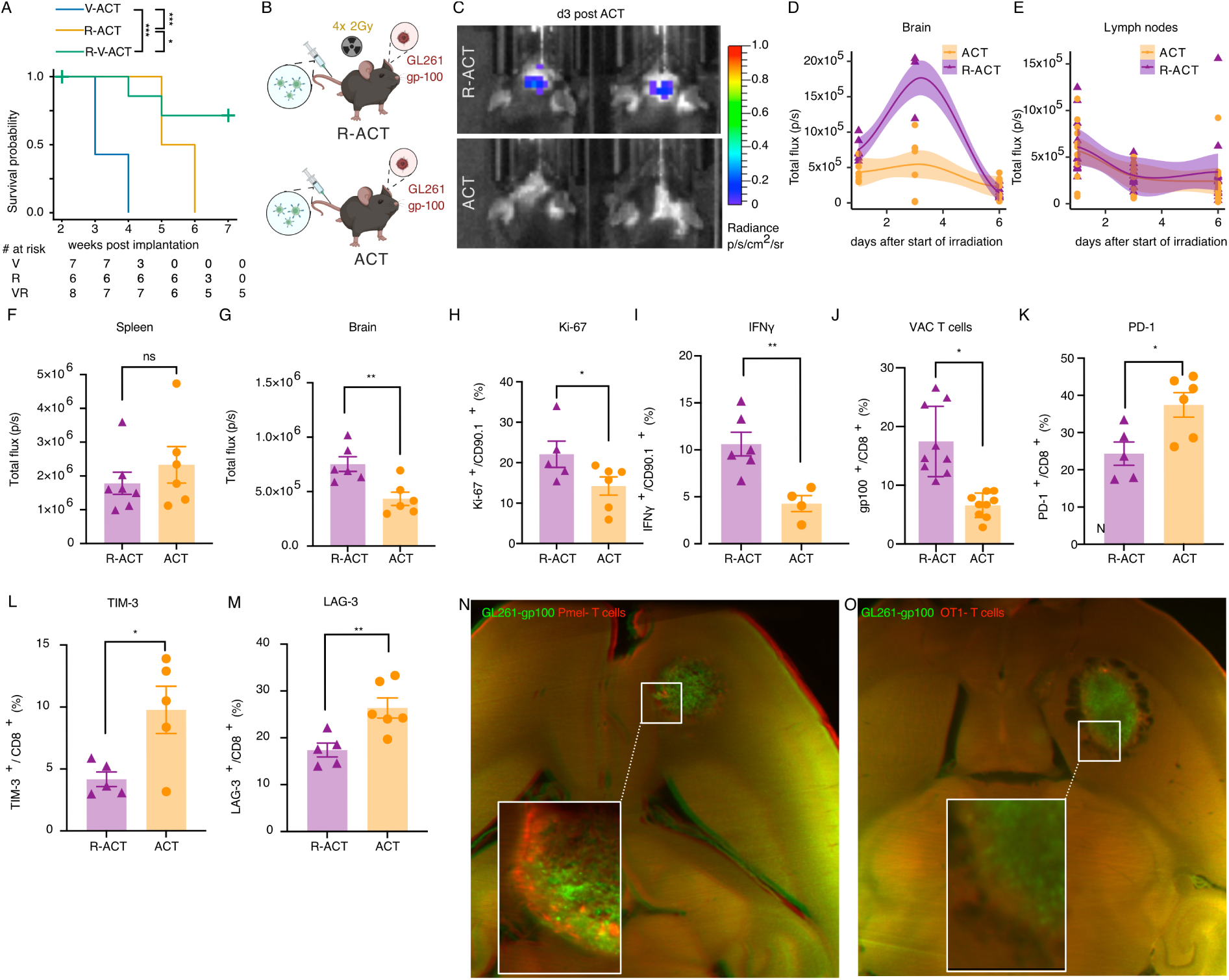
Radiotherapy enhances antigen-specific CD8⁺ T cell infiltration and survival in a gp100⁺ GL261 glioma model with adoptive T cell transfer. (A) Survival analysis showing significantly improved survival with the addition of irradiation in mice treated with adoptive cell transfer (ACT) and vaccination. V-ACT, vaccination and ACT; R-ACT, irradiation and ACT; R-V-ACT, vaccination, irradiation and ACT. (B) Treatment paradigm for (C-O). (C) Representative in vivo imaging system (IVIS) showing bioluminescence of luciferase-expressing Pmel-luc-mCherry T cells in the brain at day 3 post ACT and start of irradiation. (D-E) Quantification of bioluminescence signals from luciferase expressing T cells in the tumor (D) and lymph node (E) tracking the total influx of the antigen-specific pmel-luc-mCherry T cells. (F-M) Flow-cytometric analysis of Ki-67, IFN-γ, PD-1, LAG-3, and TIM-3 expression in tumor-infiltrating Pmel T cells nine days after ACT ± irradiation. Mean ± SEM; Student’s t-test: ** p < 0.01, * p < 0.05, ns = not significant. (N, O) Light-sheet microscopy showing gp100⁺ GFP⁺ GL261 tumor cells (green) and infiltrating Pmel-luc-mCherry T cells (red) (N) and OT1 T cells (red) (O) demonstrating specificity.

### Irradiation drives clonal expansion and metabolic reprogramming of endogenous tumor-infiltrating CD8⁺ T cells

To identify vascular mechanisms shaping anti-tumor T cell responses in the context of irradiation, we performed immunohistochemistry and paired single-cell RNA sequencing and V(D)J profiling of tumor-infiltrating CD3^+^ T cells and CD31^+^ endothelial cells in gp100-expressing GL261 intracranial tumors (GL261-gp100) (Supplementary Fig. 2A). Fractionated irradiation of established GL261-gp100 tumors resulted in a marked increase in both the proportion and absolute number of intra-tumoral CD8⁺ T cells compared to controls both in immunohistochemistry (Fig. 2A-D) and in single-cell transcriptomics (Fig. 2E-G). This effect was particularly pronounced for CD8^+^ effector memory T cells (Fig. 2F, 2H). Irradiated tumors exhibited increased T cell receptor (TCR) clonality, characterized by a higher proportion of large clonal expansions (Fig. 2I) and a reduced Shannon diversity index relative to controls (Fig. 2J). In irradiated tumors, CD8⁺ effector T cells - either proliferating, active or exhausted - harbored the most frequent TCR clonotypes. In contrast in untreated tumors, dominant clonotypes were primarily found in the active T helper and exhausted CD8^+^ T cell clusters (Fig. 2K). Gene expression profiling of glioma-infiltrating CD8^+^ T cells revealed radiation-dependent up-regulation of mitochondrial oxidative phosphorylation and respiratory chain components such as *Atp5k*, *Atp5md*, *Atp5mpl*, *Cox17*, *Cox7c*, *Ndufa1*, *Ndufa3*, and *Tomm7*, together with stress- and immune-regulatory transcripts (*Lgals1*, *Mt1*, *Ifi27l2a*, and *Tsc22d4*), suggesting enhanced mitochondrial metabolic activity and activation of stress-adaptive programs in CD8⁺ T cells after tumor irradiation (Supplementary Fig. 2B). In line, pathway enrichment analysis revealed significant enrichment of oxidative phosphorylation, mitochondrial respiration, electron transport chain assembly, mitochondrial protein import, and protein localization pathways (Supplementary Fig. 2C). This shift towards a metabolic active state in the adaptive immune compartment was accompanied by an interferon-driven response within the glioma vasculature: While pathways related to overall endothelial cell proliferation and metabolic capacity were inhibited after irradiation, induction of interferon-inducible GTPases (*Gbp2, Gbp4, Gbp5, Igtp*) and antiviral response genes (*Ifi47*) reflect stimulation of type I and II interferon responses in glioma-associated endothelial cells (Fig. 2L, 2M).

**Fig. 2:**
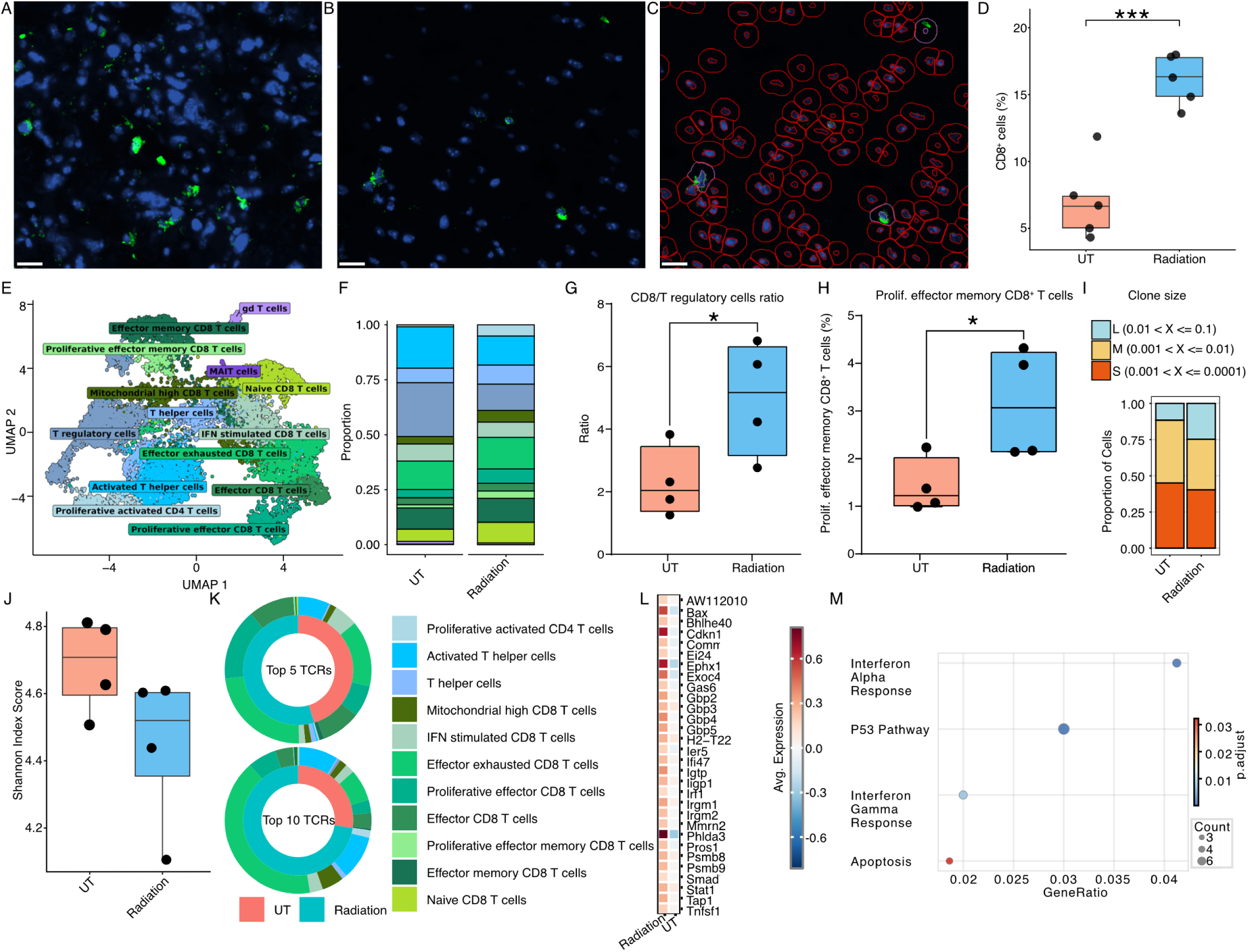
Radiotherapy increases the frequency and clonal expansion of effector CD8⁺ T cells in the tumor microenvironment of experimental GL261 gliomas. 12-week-old immunocompetent C57BL/6 mice bearing orthotopic GL261 tumors received fractionated irradiation (4x2 Gy on consecutive days); tumors were collected 24 h after the final dose for immunohistochemistry and for single-cell transcriptomic and V(D)J profiling. (A,B) CD8 (green) and DAPI (blue) staining of irradiated (A) and non-irradiated tumors (B). Scale bar, 200 μm. (C) Representative automated segmentation of non-irradiated tumor in (B) with CD8^+^ T cells segmented in purple and tumor cells in red. (D) Segmentation showed a higher percentage of CD8⁺ T cells in irradiated tumors compared with untreated (UT) controls. ***: p<0.005 (E) UMAP of single cell V(D)J data from CD45⁺ tumor-infiltrating lymphocytes isolated from irradiated (Radiation, n = 4) and non-irradiated (UT, n = 4) GL261 tumors. (F) Relative proportions of major T cell subsets. (G) CD8⁺/regulatory T cell ratio in UT versus irradiated tumors. *: p<0.05 (H) Percentage of proliferative effector memory CD8⁺ T in UT or irradiated tumors. *: p<0.05 (I) Clonal proportion analysis showing an increased fraction of medium and large clones after irradiation. (J) Shannon diversity index of TCR repertoires, showing reduced diversity and expansion of dominant clones after irradiation. (K) Phenotypic composition of the five (top) and ten (bottom) most frequent TCR clonotypes in irradiated versus control tumors. (L) Differentially expressed genes in tumor endothelial cells after irradiation; irradiated endothelial cells exhibited increased expression of interferon-stimulated and inflammatory genes (*Gbp2*, *Gbp4*, *Gbp5*, *Igtp*, *Ifi47*, and *Psmb9)* alongside stress-response and cell-cycle regulators (*Cdkn1a*, *Bax*, *Phlda3)*, consistent with interferon activation and radiation-induced endothelial reprogramming. (M) Pathway enrichment in endothelial cells from irradiated versus untreated tumors; dot size reflects gene count; color represents adjusted p value.

### Combined irradiation and vaccination synergistically promote stem-like and activated anti-tumor CD8^+^ T cell phenotypes

To dissect the impact of radiation-induced angiocrine signaling specifically on the outcome of T cell-based immunotherapies, we applied a GL261 model expressing the MHC class-I restricted ovalbumin (257-264) epitope SIINFEKL (GL261-SIINFEKL). In contrast to gp100 constituting an endogenous tumor-associated antigen, SIINFEKL is a highly immunogenic model antigen used for preclinical studies on T cell recruitment and activation (Hogquist et al., 1994). We performed single-cell RNA sequencing and V(D)J sequencing of tumor-infiltrating CD3^+^ T cells and CD31^+^ endothelial cells across four treatment conditions: adoptive transfer of *ex vivo* primed SIINFEKL-reactive (OT-I) T cells alone (ACT), ACT in combination with peptide vaccination targeting SIINFEKL (V-ACT), ACT in combination with fractionated irradiation (R-ACT) or ACT in combination with V and R (V-R-ACT) (Fig. 3A). Lung tissue served as non-CNS control to assess brain-specific T cell migration and infiltration.

**Fig. 3:**
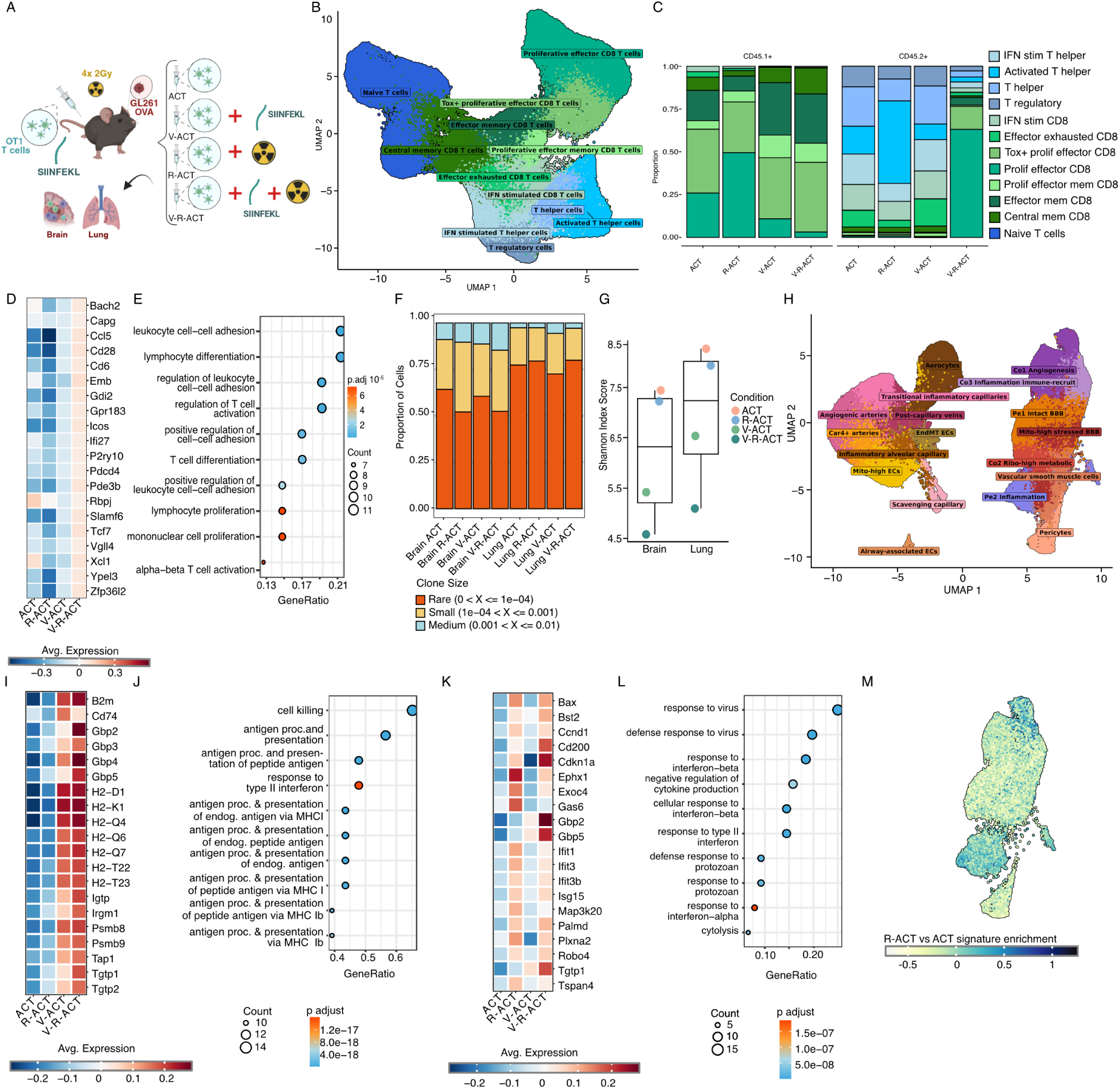
Irradiation and vaccination synergistically induce interferon-driven antigen presentation programs in tumor endothelium. (A) GL261-OVA glioma-bearing mice were treated with OT1 T cells alone (ACT) or in combination with vaccination of SINFEKL peptide (V-ACT), with 4 fractions of irradiation (R-ACT) or a triple combination of OT1 T cells, irradiation and vaccination (V-R-ACT). Lung and tumor tissue was sorted for adoptively transferred (CD45.1⁺) and endogenous (CD45.2⁺) T cells as well as endothelial cells (CD31⁺). (B) UMAP representation of T cells from all treatment groups. (C) Proportion of T cell subtypes in adoptively transferred (CD45.1⁺) and endogenous (CD45.2⁺) T cells in tumor tissue across treatment groups. (D)Top 20 upregulated genes in adoptively transferred CD8⁺ T cells in tumor tissue across treatment groups. (E) GO term enrichment in adoptively transferred CD8⁺ T cells of ACT+V+R compared to ACT+V. (F) Clonal proportion analysis of T cells demonstrating an increased proportion of medium and small clones following radiation and vaccination compared with untreated tumors and lung tissue. (G) Shannon diversity index of TCR repertoires showing reduced diversity and expansion of dominant clones after irradiation and vaccination. (H) UMAP representation of CD31⁺ endothelial cell subsets from GL261 tumor tissue across treatment conditions. (I, J) Top 20 upregulated genes in CD31⁺ endothelial cells comparing non-vaccinated and vaccinated animals across treatment conditions and corresponding Gene Ontology (GO) enrichment. (K, L) Top 20 upregulated genes in CD31⁺ endothelial cells comparing non-irradiated and irradiated tumors across treatment conditions and corresponding Gene Ontology (GO) enrichment. (M) Module score enrichment of the top 20 upregulated genes in CD31⁺ endothelial cells in irradiation signature (Fig.3.I) mapped to the brain endothelial cells UMAP.

While vaccination induced T cell memory formation, proliferating effector phenotype dominated among tumor-infiltrating OT-I CD8^+^ T cells after irradiation (Fig. 3B, C; Supplementary Fig. 3A). Combined radiation and SIINFEKL vaccination synergistically promoted an activated stem-like effector state of adoptively transferred T cells with upregulation of genes associated with T cell effector function (*Ccl5*, *Xcl1*, *Icos*, and *Slamf6*) together with interferon-alpha inducible protein 27 (*Ifi27*) and genes related to cell-to-cell adhesion (*Cd6*) (Fig. 3D). In line, gene ontology analysis of genes upregulated following combined irradiation and vaccination revealed enrichment of pathways associated with T cell activation, leukocyte adhesion, lymphocyte differentiation, and proliferative responses (Fig. 3E), consistent with the emergence of a functionally activated and immunologically engaged phenotype of OT-I CD8^+^ T cells following combination therapy.

In addition to enhancing immunological fitness of adoptively transferred tumor-reactive T cells, combined radiation and vaccination led to an increase in endogenous proliferating effector CD8^+^ T cells associated with the remodeling of the TCR repertoire: Compared to non-irradiated and non-vaccinated tumors and healthy lung tissue, V-R-ACT increased the proportion of expanded larger T cell clones, consistent with enhanced antigen-driven clonal expansion and recruitment of reactive T cell populations (Fig. 3F). Consistently, the Shannon diversity index was reduced in treated brain tissue, indicating preferential expansion of dominant clonotypes and reduced overall repertoire diversity (Fig. 3G). These findings suggest that irradiation and vaccination promote selective outgrowth of activated tumor-reactive T cell populations within the glioma microenvironment.

### Combination radioimmunotherapy induces interferon-responsive endothelial states associated with immune recruitment

To investigate the endothelial programs that may support the recruitment of the proliferative effector and effector memory T cell populations observed across treatment conditions, we next characterized the transcriptional states of glioma-associated CD31⁺ endothelial cells (Fig. 3H). Vaccination induced a distinct endothelial transcriptional program dominated by increased expression of antigen presentation and interferon-inducible genes, including *B2m, Cd74, H2-D1, H2-K1, H2-Q4/6/7, Psmb8, Psmb9*, and *Tap1* (Fig. 3I). GO analysis confirmed enrichment of pathways related to antigen processing and presentation via MHC class I and MHC Ib molecules, as well as type II interferon responses (Fig. 3J). These findings suggest that endothelial cells acquire features of immune-interacting and antigen-presenting stromal cells following vaccination, potentially reinforcing effector memory differentiation and local T cell recruitment at the tumor site.

In contrast, irradiation increased the brain-to-lung ratio of adoptively transferred OT-I cells (Supplementary Fig. 3B), suggesting that radiation promotes T cell recruitment preferentially at the level of the blood-brain barrier endothelium. Radiation induced expression of interferon-and stress-associated genes in BBB endothelial cells including *Bst2, Gbp2, Gbp5, Ifit1, Ifit3, Isg15*, and *Tgtp1*, together with genes involved in cell-cycle regulation and apoptosis such as *Cdkn1a* and *Bax* (Fig. 3K). Corresponding GO enrichment analysis demonstrated enrichment of antiviral and interferon-responsive pathways (Fig. 3L), consistent with induction of an interferon-responsive inflammatory endothelial program following irradiation.

Furthermore, vaccination-driven effector memory formation in the lung (Supplementary Fig. 3C,D) was associated with enhanced migratory capacity of OT-I cells toward the brain tumor, as assessed by a previously published brain migration signature (Kendirli et al., 2023) (Supplementary Fig. 3E) and consistent with the acquisition of an effector-like phenotype accompanied by upregulation of chemokines and adhesion molecules.

### Endothelial–T cell communication networks are remodeled by irradiation and vaccination

To further investigate treatment-associated changes in the endothelial – T cell cross-talk, we performed ligand–receptor interaction analysis between CD31⁺ tumor-associated endothelial cells and adoptively transferred CD45.1⁺ or endogenous CD45.2^+^ CD8⁺ T cells. At the global endothelial level, irradiation predominantly enhanced Notch-engagement interactions, while vaccination additionally induced endothelial immune-adhesion signatures characterized by Vcam1 and Icam1 interactions (Supplementary Fig. 4A-C).

Given that radiation effects on the vasculature are likely concentrated in specific endothelial subpopulations rather than uniformly distributed across the BBB, we next identified endothelial phenotypes previously described in human tumors (Xie et al., 2021) representing key checkpoints for T cell entry (Fig. 3H). Projection of the radiation-induced interferon gene signature (Fig.3K) onto these subpopulations revealed preferential enrichment in Pe2 inflammatory endothelial cells (Fig.3M), pointing to this subset as the principal candidate mediator of radiation-driven T cell recruitment. Indeed, radiation increased ICAM, VCAM-, and extracellular matrix-associated interactions between Pe2 inflammatory endothelial cells and both proliferative effector and effector memory CD8⁺ T cells (Fig. 4A, 4B), established drivers of T cell transmigration across the blood-brain barrier(Mapunda et al., 2022). Supporting these predicted interactions, *Icam* protein abundance was significantly higher in Pe2 inflammatory endothelial cells following irradiation (Fig. 4C).

**Fig. 4:**
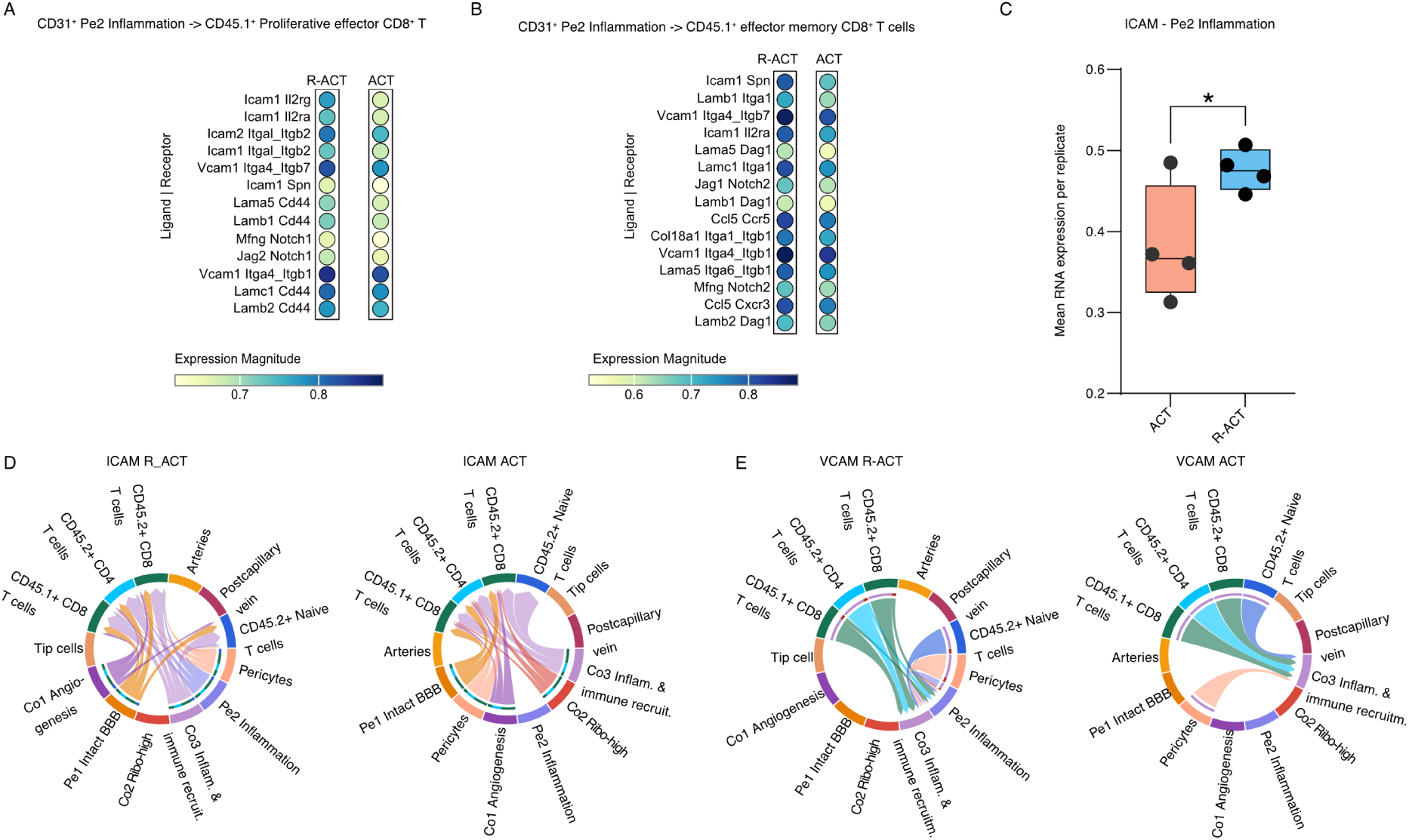
Irradiation and vaccination remodel endothelial–T cell communication through inflammatory ICAM-1- and VCAM-1-expressing endothelial subsets. (A–B) Ligand–receptor interaction analysis between Pe2 Inflammation CD31⁺ endothelial cells and proliferative effector CD8⁺ T cells (A) and effector memory CD8⁺ T cells (B) comparing irradiated ACT-treated tumors (R-ACT) versus ACT alone. Irradiation enhances ICAM-, VCAM- and Notch-interactions between inflammatory endothelial cells and tumor-reactive T cells. (C) ICAM RNA expression in Pe2 Inflammation CD31⁺ endothelial cells. (D, E) Global cell–cell communication networks for ICAM- and VCAM-associated signaling pathways across endothelial and immune cell populations under treatment conditions. Chord diagrams illustrate predicted interactions between endothelial subsets, pericytes, angiogenic endothelial cells, inflammatory endothelial populations, and T cell populations within the glioma microenvironment.

In addition, Pe2 endothelial cells displayed enriched Notch pathway interactions with CD8⁺ T cells, including *Dll4-*, *Jag2*-, and *Mfng*-associated signaling through Notch1 and Notch2 receptors, particularly within the effector-memory compartment. Given the established role of Notch signaling in CD8⁺ T cell effector differentiation and memory formation (Backer et al., 2014; Kondo et al., 2017), these findings suggest that inflammatory endothelial cells contribute not only to T cell recruitment but also to the differentiation and maintenance of activated CD8⁺ T cell states within irradiated tumors.

Global interaction network analysis further demonstrated increased connectivity involving inflammatory (Pe2) and immune-recruiting (Co3) endothelial subsets in irradiated and vaccinated gliomas (Figure 4D, E), consistent with therapy-induced vascular remodeling that promotes immune recruitment and endothelial-T cell crosstalk within the glioma microenvironment.

### Irradiation-induced endothelial–T cell interactions are recapitulated in human glioblastoma

To determine whether the irradiation-driven endothelial–T cell interactions identified in murine models are recapitulated in human glioblastoma (GBM), we analyzed a cohort of nine patients with isocitrate dehydrogenase (IDH)-wildtype GBM who had undergone surgical resection followed by radiotherapy and temozolomide chemotherapy (Seferbekova et al., 2026). Among these, four patients exhibited histologically confirmed tumor recurrence (progressive disease; PP), four showed no evidence of recurrence but displayed radionecrotic changes (radiation injury; RI), and one patient displayed histopathological features of both conditions.

Spatial profiling revealed enrichment of T cells within perivascular niches in both PP and RI samples (Fig. 5A; Supplementary Fig. 5A, B), indicating that vascular-associated immune infiltration is a common feature following treatment. However, quantitative analysis demonstrated a significantly higher accumulation of both CD4⁺ and CD8⁺ T cells in close proximity to blood vessels in RI samples compared to PP samples (Fig. 5B, C). This suggests that enhanced perivascular T cell infiltration is associated with, and may contribute to, the development of radionecrosis following irradiation. In contrast, no comparable increase was observed for myeloid cell populations (Fig. 5D), indicating a degree of specificity in the immune response.

**Fig. 5:**
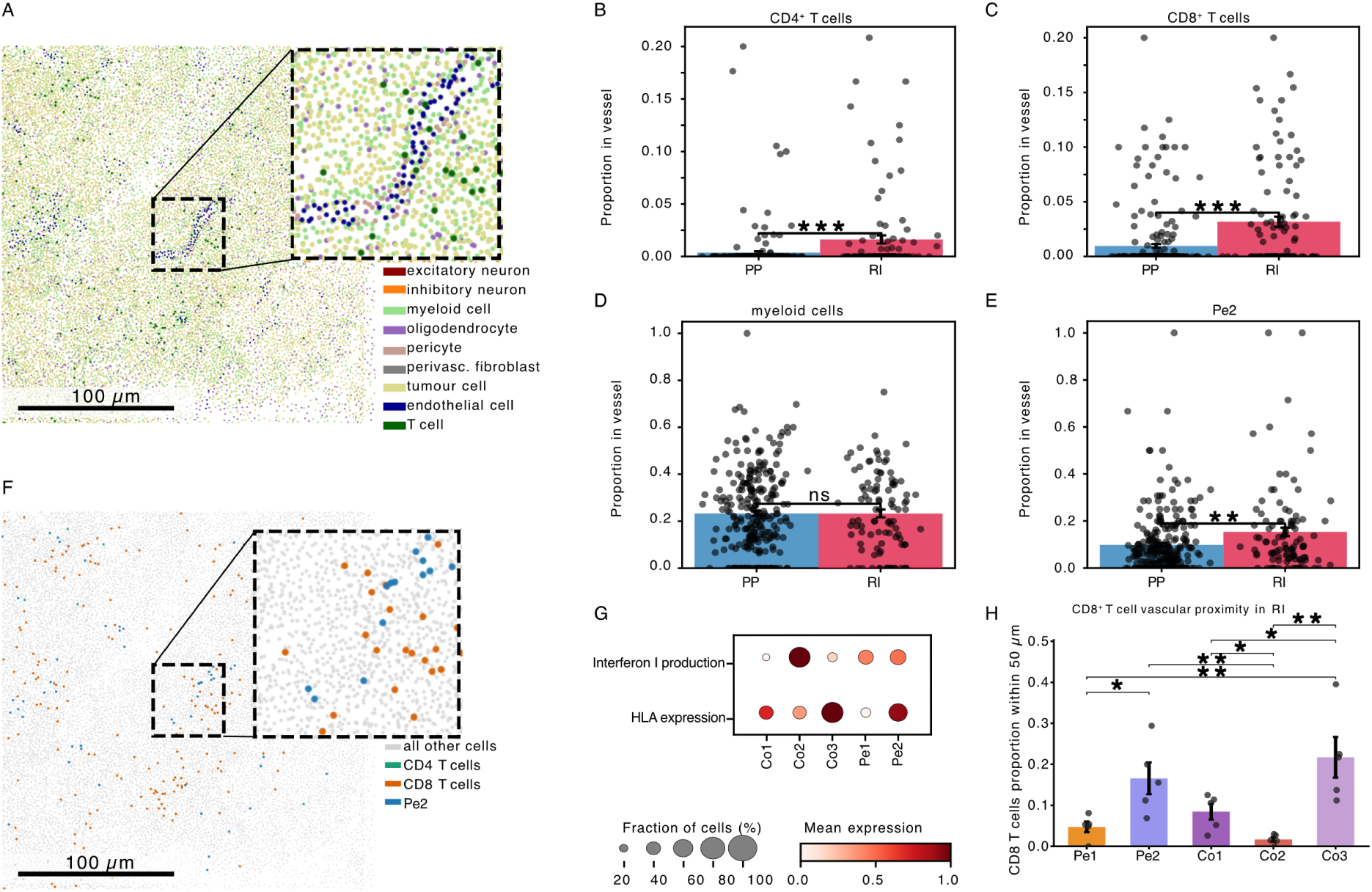
Perivascular enrichment of T cells and association with the Pe2 endothelial subset in irradiated human glioblastoma. (A) Spatial transcriptomics map of a representative recurrent glioblastoma section showing the distribution of major cell types, including endothelial cells and T cells, with a zoom-in highlighting perivascular niches. (B–C) Quantification of CD4⁺ (B) and CD8⁺ (C) T cells located within vessel-associated regions, demonstrating a significantly higher proportion of both subsets in radiation injury (RI) compared to progressive disease (PP). Each point represents an individual observation; bars indicate mean ± s.d.; ***P < 0.001. (D) Proportion of myeloid cells associated with vessels in PP and RI samples. (E) Proportion of endothelial cells belonging to the Pe2 subset in vessel-associated regions, revealing a significant enrichment in RI compared to PP. (F) Dot plot summarising expression of antigen presentation (HLA) and interferon response (type I interferon production) signatures across endothelial subtypes, highlighting elevated interferon-related signalling and antigen presentation features within specific endothelial populations, including Pe2. (G) Spatial map highlighting CD4 ⁺ T cells, CD8⁺ T cells, and Pe2 endothelial cells, illustrating their co-localisation within perivascular niches. (H) Quantification of CD8⁺ T cells located within endothelial subsets vessel-associated regions (50uM) in radiation injury (RI). Each point represents patient; bars indicate mean ± s.d.; ***P < 0.001.

Notably, the increased T cell infiltration in RI samples coincided with a significantly higher proportion of endothelial cells belonging to the Pe2 subset (Fig. 5E, F), whereas no such enrichment was observed for other endothelial subpopulations (Supplementary Fig. 5C, D). In agreement with the interferon-responsive endothelial state observed in irradiated murine gliomas, Pe2 endothelial cells displayed high interferon production-related gene scores, while both Pe2 and Co3 populations were enriched for HLA expression, indicative of enhanced immunoregulatory and antigen-presenting capacity (Fig. 5G). Spatial mapping of CD8^+^ T cell abundance across endothelial subsets further confirmed that Pe2 and Co3 endothelial cells displayed the highest density of perivascular CD8^+^ T cell clustering (Fig. 5H). Collectively, these findings support the notion that the Pe2 endothelial subset plays a conserved role in mediating T cell recruitment to the perivascular niche following irradiation in human GBM.

## Discussion

Glioblastoma remains a profoundly immune-excluded tumor in which therapeutic efficacy is limited not only by immunosuppressive signaling but also by restricted immune cell access to the central nervous system. In this study, we identify radiation-induced endothelial activation as a key mechanism that enables the recruitment and local expansion of tumor-specific CD8⁺ T cells within the glioma microenvironment. These findings extend current models of radiotherapy-induced immune modulation by positioning the tumor vasculature as an active regulator of anti-tumor immunity in the brain.

Consistent with prior reports linking radiotherapy to enhanced anti-tumor immune responses (Burnette et al., 2011; Vanpouille-Box et al., 2017), we observed increased accumulation and clonal expansion of tumor-specific CD8⁺ T cells following irradiation, accompanied by enrichment of proliferative, cytotoxic, and interferon-responsive effector states. These effects were observed across both an endogenous tumor antigen model (gp100) and an ovalbumin (OVA) model antigen system, supporting the generalizability of radiation-enhanced T cell responses beyond a single antigenic context. Notably, this response was largely confined to the tumor site, supporting emerging evidence that local proliferation and tissue adaptation—rather than systemic expansion alone—are critical determinants of effective T cell immunity in solid tumors, including glioma (Watowich et al., 2023; Yost et al., 2019). The observed increase in TCR clonality further suggests selective expansion of tumor-reactive clones, a feature associated with effective anti-tumor responses (Brastianos et al., 2017; Wu et al., 2020) and increasingly recognized as a hallmark of locally expanding, tissue-adapted T cells in glioma (Kilian et al., 2022; Watowich et al., 2023).

A central finding of our study is that radiation reprograms the glioma vasculature into an interferon-responsive, immune-permissive niche that facilitates T cell recruitment. While radiation-induced interferon signaling has been extensively characterized in tumor and immune cells, our data demonstrate that endothelial cells undergo a coordinated interferon-driven reprogramming that includes upregulation of antigen presentation machinery and adhesion molecules such as ICAM-1 and VCAM-1. These changes are functionally positioned to enhance multiple steps of leukocyte trafficking across the blood–brain barrier. This finding supports a model in which vascular regulation of immune cell trafficking—long recognized as a key constraint in the central nervous system—is dynamically modulated by irradiation, converting the typically restrictive blood–brain tumor barrier into a more immune-permissive interface (Arvanitis et al., 2020).

Our findings extend earlier observations that radiation profoundly alters endothelial biology by inducing DNA damage responses, inflammatory signaling, vascular permeability changes, senescence, and adhesion molecule expression (Hallahan et al., 1996). In peripheral tumor models, radiation-induced endothelial activation has been linked to enhanced immune cell recruitment and improved T cell access to tumors (Ganss et al., 2002). Consistent with these observations, endothelial activation through ICAM-1 and VCAM-1 upregulation is a prerequisite for leukocyte adhesion and transmigration across the blood-brain barrier (Mapunda et al., 2022). Previous studies in syngeneic intracranial glioma models further demonstrated that combining whole-brain irradiation with peripheral immunotherapeutic strategies, including GM-CSF–secreting tumor vaccines or agonistic anti-CD137 antibodies, resulted in durable tumor control and long-term survival associated with increased tumor-infiltrating lymphocytes and enhanced tumor-specific IFN-γ production (Newcomb et al., 2006). Our data extend these findings by identifying radiation-induced endothelial reprogramming as a potential mechanistic link between irradiation and enhanced recruitment of tumor-reactive CD8⁺ T cells within the glioma microenvironment.

Mechanistically, the endothelial response we describe is consistent with activation of type I interferon pathways downstream of cytosolic DNA sensing, such as cGAS–STING signaling (Deng et al., 2014; Vanpouille-Box et al., 2017). While this pathway has primarily been studied in dendritic cells as a driver of T cell priming, our findings suggest that it simultaneously licenses the tumor vasculature to support T cell entry. This coordinated activation of immune and vascular compartments may represent a critical requirement for effective anti-tumor immunity in glioblastoma, where failure of immune cell trafficking constitutes a major therapeutic barrier (Bunse et al., 2025). Although our data strongly associate endothelial interferon activation with enhanced T cell recruitment, future studies employing endothelial-specific perturbation strategies will be required to establish direct causality.

Importantly, we demonstrate that these vascular changes have direct functional consequences for immunotherapy. Radiotherapy significantly enhanced the efficacy of both adoptive T cell transfer and antigen-specific vaccination, resulting in increased brain-specific T cell accumulation, proliferation, and effector differentiation. These findings support a model in which radiotherapy acts not only as an immune adjuvant but also as a facilitator of immune cell access to the tumor site. In this framework, radiation-induced endothelial activation and antigen-specific T cell priming represent complementary processes that together enable effective anti-tumor responses. This concept is consistent with prior observations of synergy between radiotherapy and immunotherapy (Demaria et al., 2015; Zeng et al., 2013), but provides a mechanistic basis rooted in vascular–immune crosstalk.

From a translational perspective, our findings have several implications for the design of combination therapies in glioblastoma. First, they suggest that radiotherapy may be a critical prerequisite for effective T cell–based therapies, including adoptive cell transfer and neoantigen vaccination approaches currently under clinical investigation (Brown et al., 2024; Bunse et al., 2026; Grassl et al., 2023; Majzner et al., 2022). By enhancing endothelial activation and adhesion molecule expression, radiation may overcome one of the principal barriers to immunotherapy in the brain — insufficient T cell trafficking. Second, the identification of ICAM-1 and VCAM-1 – mediated interactions as key components of this process raise the possibility that endothelial activation could serve as a biomarker of therapeutic responsiveness or as a target for pharmacologic modulation. Third, our data highlight the importance of treatment sequence, as the transient nature of radiation-induced interferon signaling and endothelial activation may define a therapeutic window during which T cell recruitment is maximized.

While our data implicate endothelial activation as a key regulator of T cell recruitment, direct causal validation — for example through endothelial-specific perturbation of adhesion molecules or interferon signaling pathways — will be important to further delineate these mechanisms. In addition, radiation-induced interferon responses are likely to be context-dependent and may also promote adaptive resistance, including upregulation of immune checkpoint pathways (Dovedi et al., 2014), supporting the rationale for combination strategies incorporating checkpoint blockade.

In conclusion, our findings identify radiation-induced endothelial activation as a key mechanism shaping T cell trafficking and function in murine glioblastoma models. By demonstrating that radiotherapy reprograms the tumor vasculature into an interferon-driven, immune-permissive state, this work provides a mechanistic framework for understanding how irradiation can facilitate T cell access to brain tumors. Given that radiotherapy is already a standard component of glioblastoma treatment, these results support its rational integration with immunotherapeutic strategies, particularly vaccination and adoptive cell transfer approaches, in which efficient T cell recruitment to the tumor may be critical for therapeutic efficacy.

---

**Supplementary Fig. 1:**
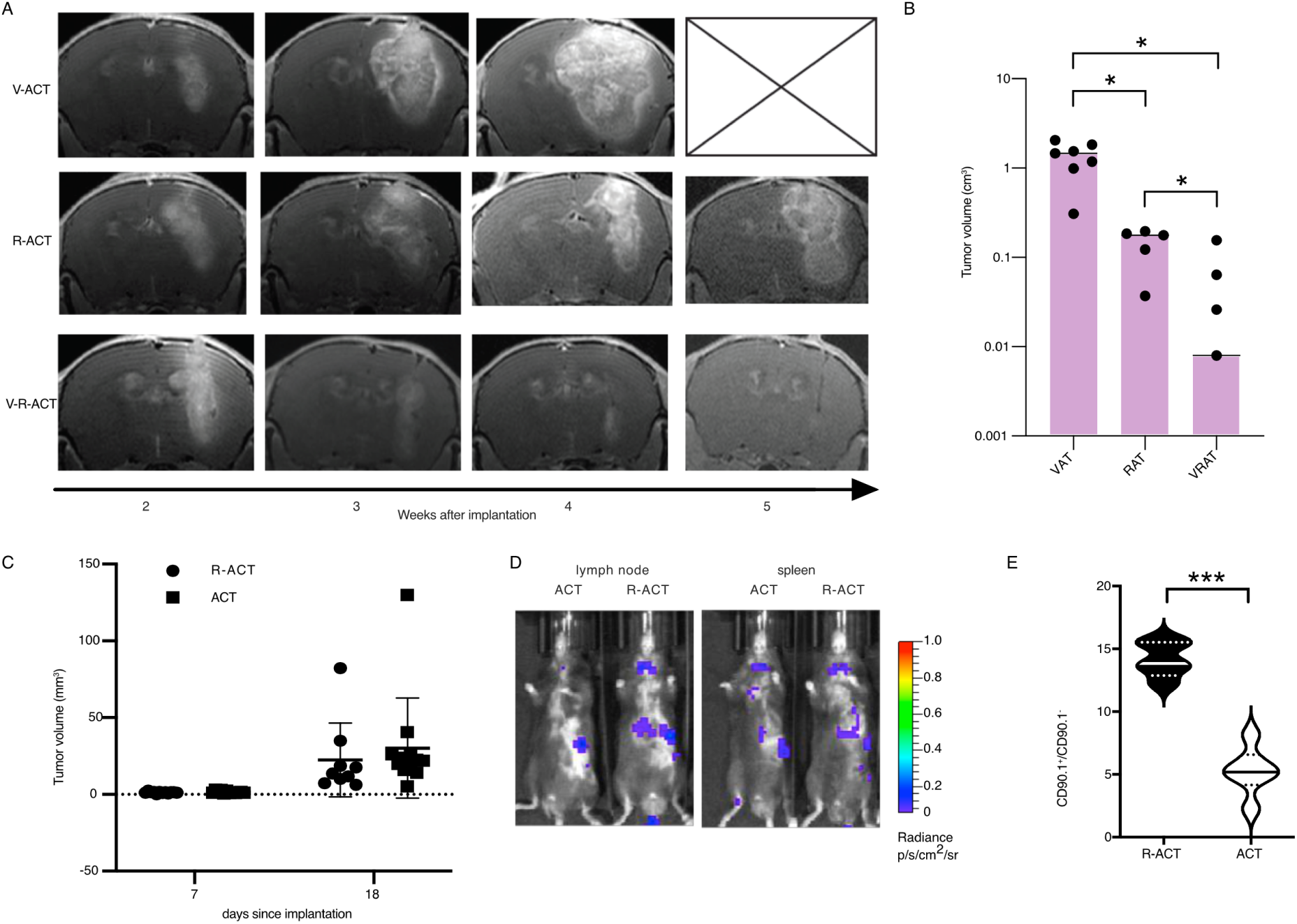
Radiotherapy enhances tumour control and promotes antigen-specific T cell accumulation following adoptive transfer. (A) Representative T1w, Gd-enhanced MRI images of GL261 gp100⁺ tumours at indicated time points following treatment with vaccination and adoptive T cell transfer (VAT), irradiation plus adoptive transfer (RAT), or the combination of vaccination, irradiation, and adoptive transfer (VRAT), showing reduced tumour burden in combined treatment groups over time. (B) Quantification of tumour volumes across treatment groups at week 4 after implantation demonstrating significantly reduced tumour size upon combination therapy (VRAT) compared to VAT or RAT alone. (C) Longitudinal measurement of tumour volume following tumor implantation prior to adoptive transfer (AT) and irradiation showing no significant difference in tumor volumes prior to treatment initiation between groups. (D) Bioluminescence imaging of lymph nodes and spleen demonstrating no significant difference in systemic distribution of transferred T cells in irradiated mice (RAT) relative to ACT controls. (E) Quantification of CD90.1⁺ adoptively transferred T cells relative to endogenous CD90.1⁻ cells, showing increased frequency of transferred T cells in irradiated conditions. Data are presented as mean ± SEM; statistical significance determined by Student’s t-test (*P < 0.05, ***P < 0.001).

**Supplementary Fig. 2:**
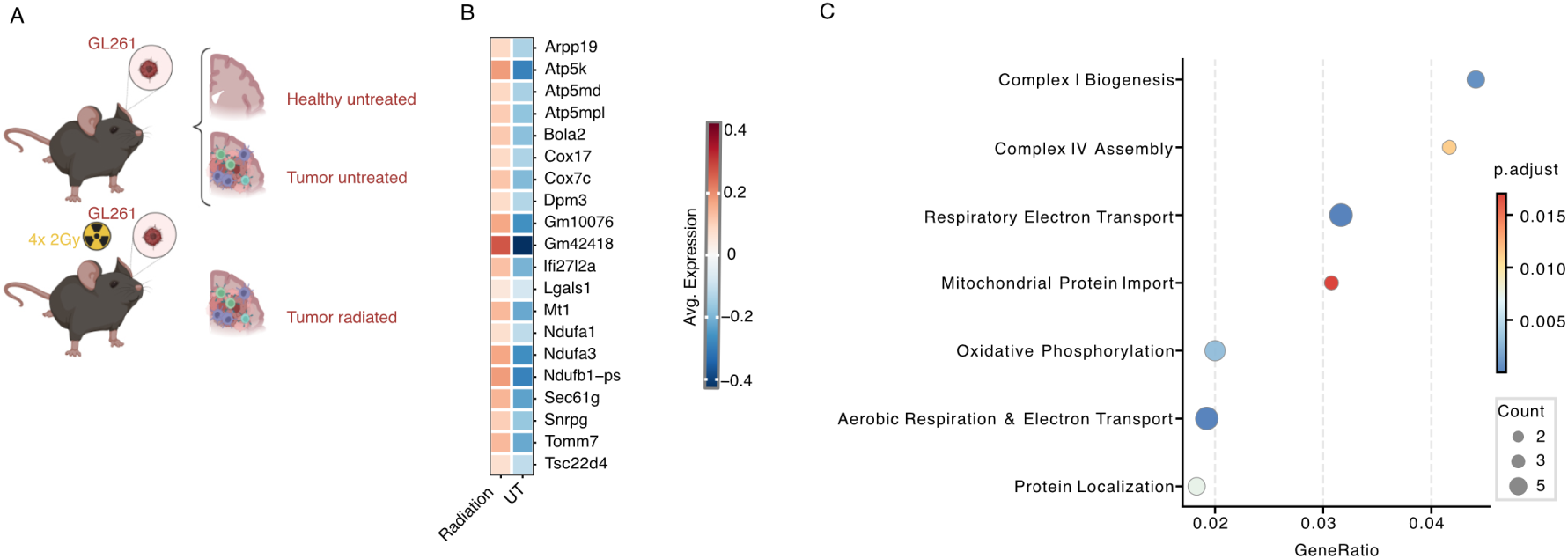
Irradiation promotes T cell infiltration and mitochondrial metabolic remodeling in CD8⁺ effector T cells. (A) Experimental workflow: 12-week-old immunocompetent C57BL/6 mice bearing orthotopic GL261 tumors were treated with a fractionated dose of 4x2 Gy irradiation on consecutive days; tumors were collected 24 h later for single cell transcriptomic and V(D)J profiling. (B) Heatmap of genes upregulated in CD8⁺ T cells, highlighting enrichment of mitochondrial oxidative phosphorylation and respiratory chain transcripts (Atp5k, Atp5md, Ndufa1, Cox17, Tomm7) together with stress- and immune-regulatory genes including Lgals1, Mt1, and Ifi27l2a. (C) Combined pathway enrichment analysis of differentially expressed genes in CD8⁺ T cells from irradiated versus untreated tumors. Dot plot showing enrichment of oxidative phosphorylation, mitochondrial respiration, electron transport chain assembly, and protein localization pathways. Dot size indicates gene count and color represents adjusted p value.

**Supplementary Fig. 3:**
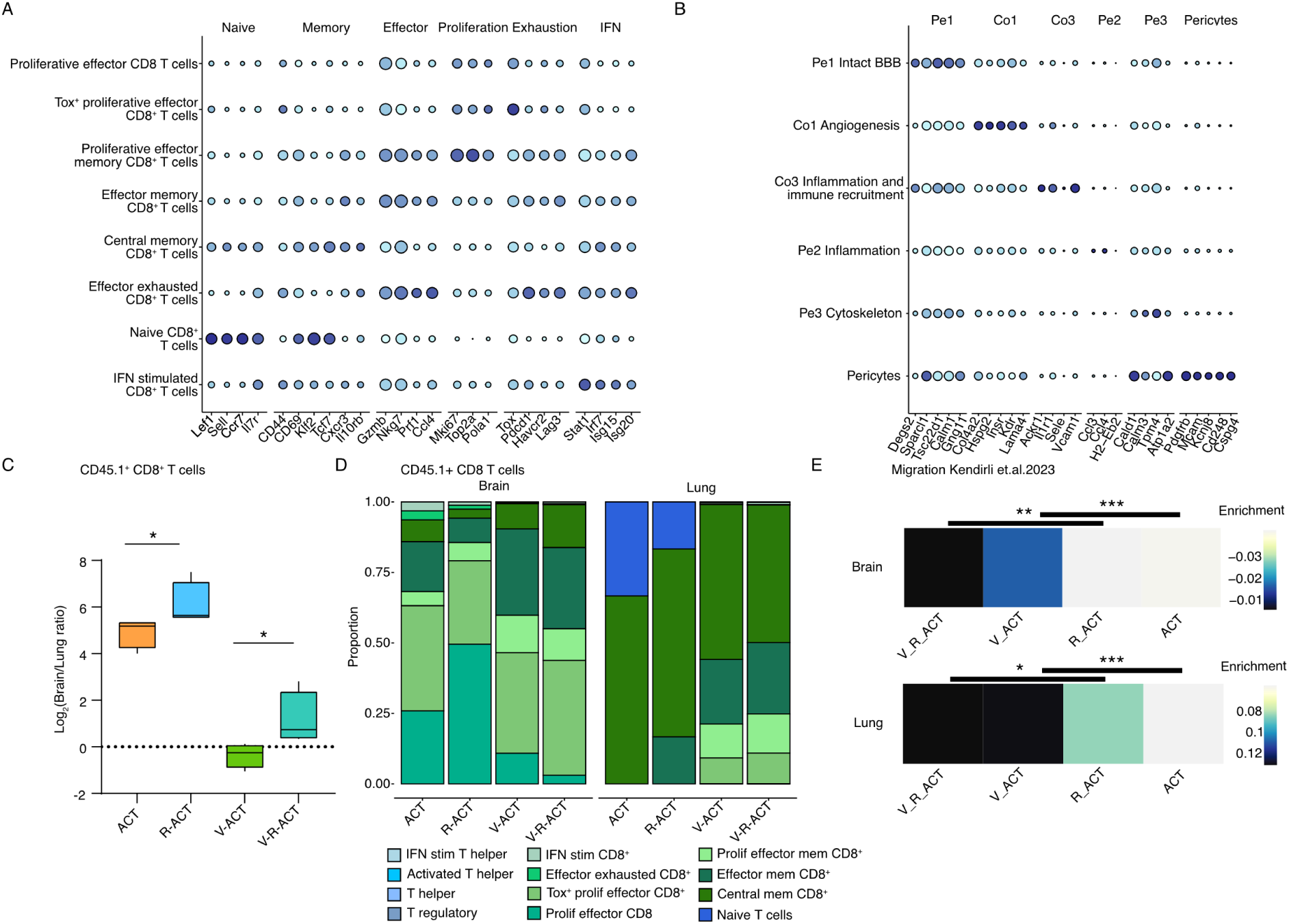
Subset marker expression in GL261-OVA glioma-bearing mice. (A) Proportion of adoptively transferred (CD45.1^+^) T cell subtypes in brain and lung across treatment groups. (B, C) Dot plot showing expression of canonical marker genes across T cell (B) as well as endothelial and perivascular cell (C) subsets identified by single-cell RNA sequencing. Dot size represents the proportion of cells expressing a given gene and color indicates average expression level. (D) Log2 brain to lung ratios of adoptively transferred (CD45.1^+^) CD8^+^ T cells in each of the therapy conditions. (E) Module score enrichment of the T cell brain migration signature derived from the CRISPR screen reported in(Kendirli et al., 2023).

**Supplementary Fig. 4:**
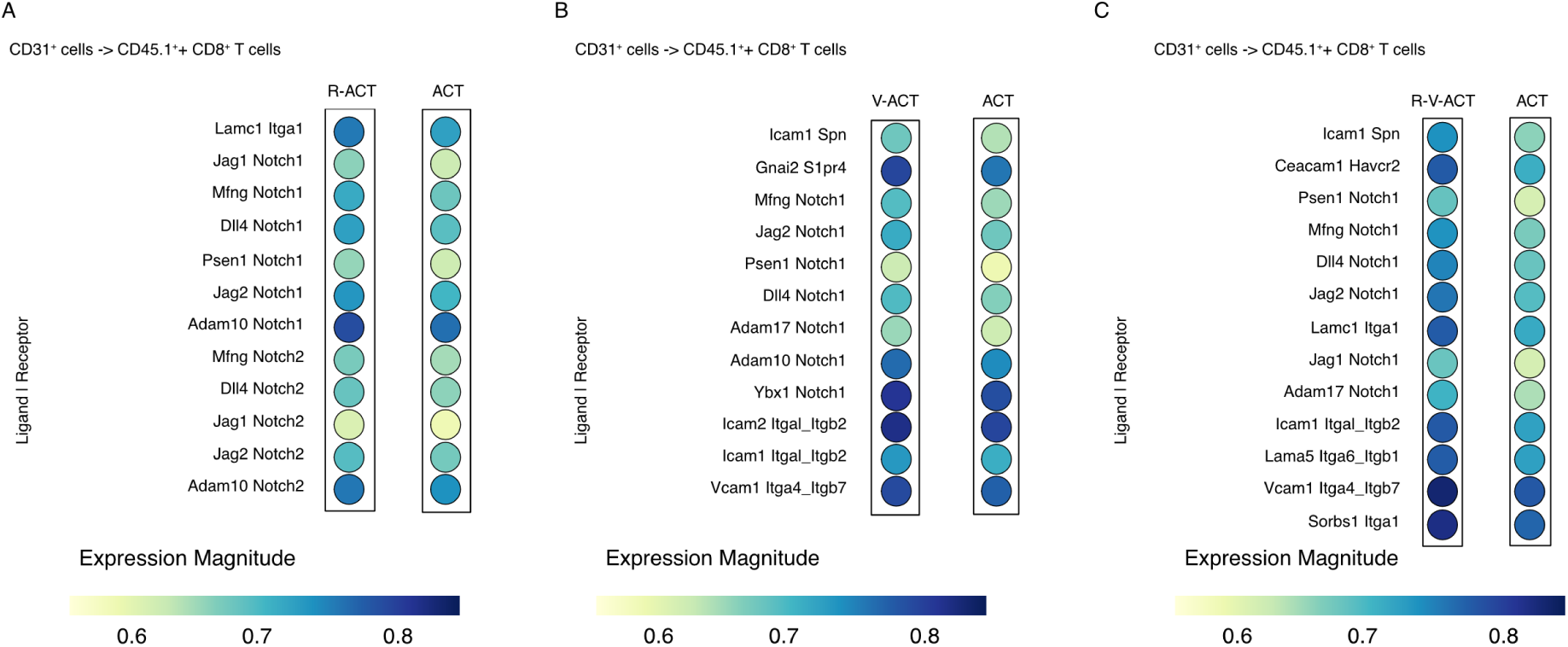
Irradiation and vaccination remodel endothelial–T cell communication networks in the glioma microenvironment. (A–C) Unbiased ligand–receptor interaction analysis between CD31⁺ endothelial cells and adoptively transferred CD45.1⁺ CD8⁺ T cells across treatment conditions. (A) Comparison of irradiated ACT-treated tumors (R-ACT) versus ACT alone reveals enrichment of endothelial Notch-associated signaling pathways, including Jag1/2–Notch1/2, Dll4–Notch1/2, Adam10–Notch1/2, and Mfng–Notch1/2 interactions, together with extracellular matrix-associated signaling. (B) Comparison of vaccinated ACT-treated tumors (V-ACT) versus ACT demonstrates increased adhesion- and trafficking-associated interactions, including ICAM- and VCAM-dependent signaling pathways. (C) Comparison of combined vaccination and irradiation (V-R-ACT) versus ACT shows further enrichment of endothelial adhesion, extracellular matrix, and immunoregulatory signaling pathways, including CEACAM1–HAVCR2 interactions. Together, these analyses identify endothelial adhesion and trafficking programs as major features of therapy-induced vascular–immune crosstalk in glioma.

**Supplementary Fig. 5:**
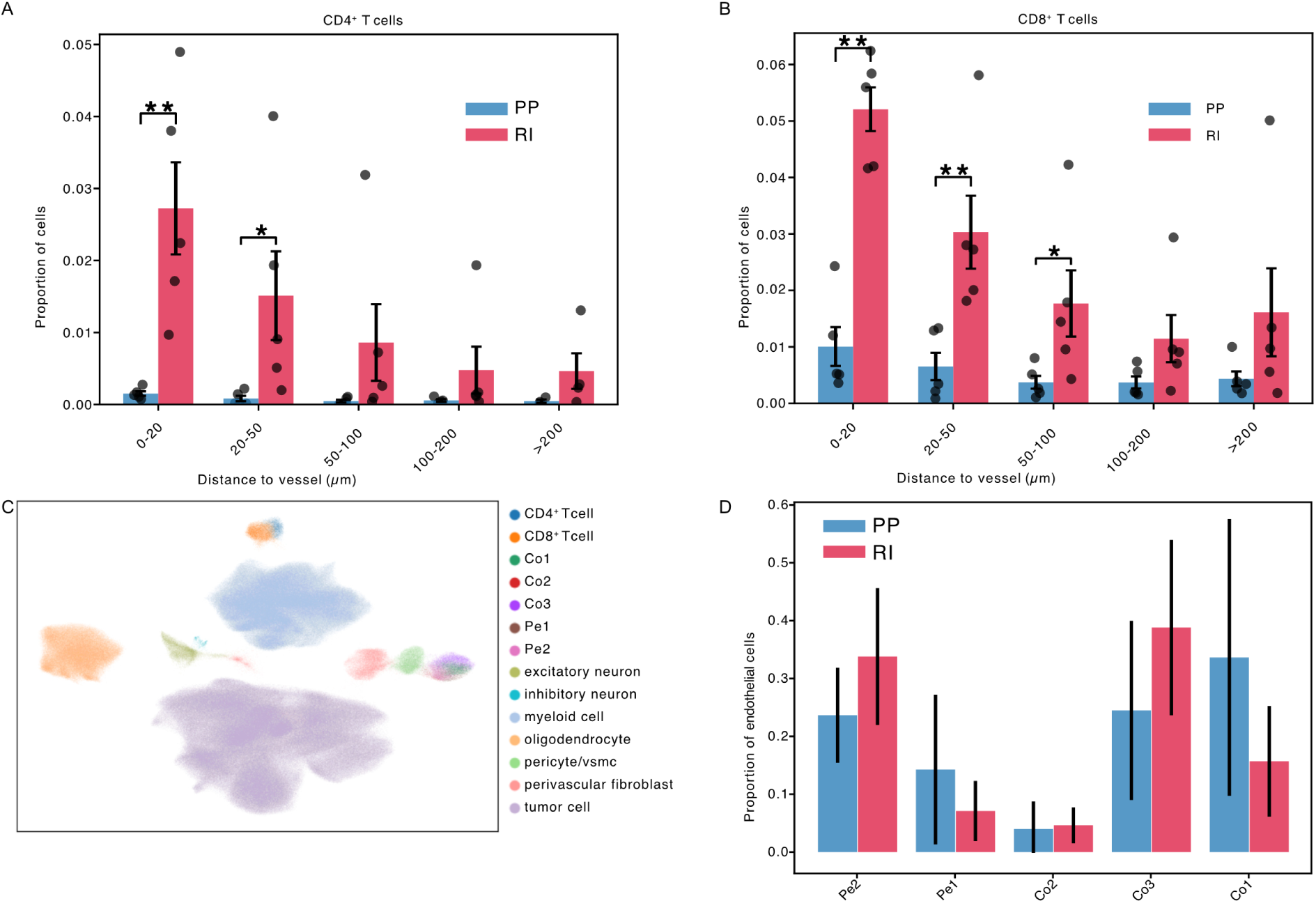
Spatial distribution of T cells and endothelial cell subtype composition in glioblastoma samples. (A–B) Quantification of CD4⁺ (A) and CD8⁺ (B) T cells as a function of distance to blood vessels in progressive disease (PP) and radiation injury (RI) samples. Both CD4⁺ and CD8⁺ T cells are enriched in close proximity to vessels (0–20 µm), with significantly higher proportions observed in RI compared to PP, particularly within perivascular regions. Data are shown as mean ± s.d. (*P < 0.05, **P < 0.01). (C) UMAP embedding of single-cell transcriptomic data showing major cell populations, including endothelial subtypes (Co1–3, Pe1–2), T cells, myeloid cells, and neural and stromal populations. (D) Proportional distribution of endothelial cell subtypes in PP and RI samples, highlighting an increased contribution of the Pe2 subset in RI relative to PP, while other endothelial populations show no consistent enrichment.

## Materials and Methods

### Study design

The aim of this study was to investigate how radiotherapy reshapes endothelial–immune interactions within the glioma microenvironment and how these changes influence T cell trafficking and immunotherapy efficacy. Orthotopic murine glioma models expressing either gp100 or SIINFEKL were combined with adoptive T cell transfer, peptide vaccination, and fractionated irradiation. Tumor growth, immune infiltration, endothelial phenotypes, and T cell clonality were analyzed using flow cytometry, immunohistochemistry, bioluminescence imaging, single-cell RNA sequencing (scRNA-seq), V(D)J sequencing, and ligand–receptor interaction analyses. Human spatial transcriptomic datasets were analyzed for validation of radiation-associated endothelial–immune phenotypes.

### Cell lines

Gl261 cells were obtained from the Division of Cancer Treatment and Diagnosis at the National Cancer Institute. GL261^gp100^ and GL216^OVAsiinfekl^ cell lines were generated by transfecting GL261 cells with the pMXS-gp100-IRES-blasticidine or pMXS-OVA_siinfekel_-IRES-blasticidine plasmids. Transfected cells were selected using 9 μg/mL blasticidin (Gibco). Cells were maintained in Dulbecco’s Modified Eagle’s Medium (DMEM) supplemented with 10% fetal bovine serum (FBS) and 1% penicillin–streptomycin. All cultures were maintained at 37 °C in a humidified incubator with 5% CO₂, and the cell lines were routinely screened for mycoplasma contamination.

### Mice

C57BL/6J mice were purchased from the Jackson Laboratory and bred at the DKFZ animal facility. PMEL mice harbor a rearranged transgenic T cell receptor that specifically recognizes pmel-17, the murine homolog of the human SILV antigen (gp100), were used as donors of primary murine T cells for experiments in the gp100 tumor model. OT-I mice express a transgenic T cell receptor specific for ovalbumin residues 257–264 and were used as a source of T cells for studies in the OVA model. OT-I and Pmel-1 mice were bred at the DKFZ animal facility. Mice were housed under Specific and Opportunistic Pathogen Free (SOPF) conditions and under 12-h day/night cycle. Mice were between 8-10 weeks at the time of the surgery and were assigned to age-matched and sex-matched experimental groups. All animal procedures followed the institutional laboratory animal research guidelines and were approved by the governmental authorities (Regional Administrative Authority Karlsruhe, Germany. Animal approval protocols are G27-17, G263/18, G264/18 and G-170/21.

### Intracranial surgery for tumor inoculation

GL261^gp100^ and GL216^OVAsiinfekl^ cells were harvested in PBS and stereotactically implanted into the right hemisphere at a density of 1 × 10⁵ cells per 2 μL PBS per mouse. With a 10 μL Hamilton micro-syringe driven by a fine step stereotactic device (Stoelting), the tumor cells were inoculated 2 mm right lateral of the bregma and 1 mm anterior to the coronal suture with an injection depth of 3 mm below the dural surface. The surgery was conducted under anesthesia using ketamine (100 mg/kg, intraperitoneal) and xylazine (10 mg/kg, intraperitoneal). Following the procedure, mice were provided with analgesic treatment for three days. Animals were monitored daily for tumor-associated symptoms, and euthanasia was performed once predetermined endpoint criteria were reached or if neurological deficits were observed.

### Irradiation

Mice were anesthetized with a fully reversible injection consisting of medetomidine (0.5 mg/kg) and midazolam (5 mg/kg), administered subcutaneously as a combined injection (total volume ≤ 0.1 ml/10 g body weight, dissolved in sterile 0.9% NaCl). Anesthetic depth was confirmed by absence of hind-limb withdrawal reflexes. Antagonization was performed immediately after irradiation using atipamezole (2.5 mg/kg) and flumazenil (0.5 mg/kg), administered subcutaneously into the neck as a combined injection (total volume 0.1 ml/20 g body weight, dissolved in sterile 0.9% NaCl). Control animals that did not receive irradiation underwent the same anesthesia and antagonization protocol (mock treatment).

Right-hemisphere irradiation was performed using therapeutic local photon irradiation at a dose of 2 Gy per session using MultiRad25 on four consecutive days. The remaining body was shielded from radiation using lead shielding.

### Adoptive transfer of CD8^+^ OT-I and Pmel T cells

CD8^+^ T cells were isolated from OT-I mice for the GL216^OVAsiinfekl^ model and Pmel T cell for the GL261^gp100^ model. For the bioluminescence experiment Pmel-Liceferase+ T cells were adoptively transferred into GL261^gp100^ tumor-bearing mice.

Mice were anesthetized and perfused with 1x PBS. Spleen and lymph nodes were extracted and meshed through 100/70 μm smart-strainers. Red blood cells (RBCs) were removed using red cell lysis solution (Miltenyi). T Cells were isolated using negative selection by CD8^+^ isolation kit (Miltenyi). Mice were briefly warmed under a red-light lamp prior to intravenous injection of 5 × 10^6^ cells in 150 μL PBS per mouse into the tail vein of GL216^OVAsiinfekl^ bearing mice at day 4 of tumor irradiation. To distinguish transferred and endogenous T cells, congenic markers CD45.1 and CD45.2 were used. Transferred OT-I T cells were identified as CD45.1^+^ cells, whereas endogenous host-derived T cells were identified as CD45.2^+^ cells.

### Vaccine

Either gp100 or ovalbumin (OVA) peptides were administered intravenously (i.v.) to GL261^gp100^ and GL216^OVAsiinfekl^ tumor-bearing mice, respectively. Each mouse received a dose of 50 μg peptide in 100 μL PBS.

### MRI and response criteria

Magnetic resonance imaging (MRI) was performed at the Small Animal Imaging Core Facility of the DKFZ using a Bruker BioSpec 3 Tesla system (Ettlingen, Germany) equipped with ParaVision software (version 360 V1.1). Mice were anesthetized with 3.5% sevoflurane delivered in air during imaging. Lesions were identified using T2-weighted imaging acquired with a T2_TurboRARE sequence with the following parameters: TE = 48 ms, TR = 3350 ms, field of view (FOV) 20 × 20 mm, slice thickness 1.0 mm, 3 averages, scan time 3 min 21 s, echo spacing 12 ms, RARE factor 8, 20 slices, and an image matrix of 192 × 192. Tumor volumes were calculated through manual segmentation using Bruker ParaVision software version 6.0.1.

Mice that did not develop tumors were terminated by cervical dislocation after day 20.

### Bioluminescence Imaging

Adoptively transferred luciferase-expressing T cells were tracked non-invasively using an In Vivo Imaging System (IVIS) Lumina Series III (PerkinElmer). Prior to imaging, hair was shaved from the regions of interest. Mice received a subcutaneous injection of 50 mg/kg D-Luciferin (StayBrite™, BioVision, Mountain View, CA) and anesthesia was induced with 3-4% isoflurane. Bioluminescence images were acquired 10 minutes after D-Luciferin injection at exposure times of 30, 45, and 60 seconds, while mice were maintained under 1.5% isoflurane throughout image acquisition.

Image acquisition and analysis were performed using Living Image software version 4.3 (PerkinElmer). For signal quantification, regions of interest (ROIs) were manually defined around the relevant anatomical areas, and bioluminescent signal was expressed as total photon flux (photons/second). To ensure comparability across time points, identical ROIs were applied consistently throughout each experiment.

### Light-sheet fluorescence microscopy

Whole brains were processed for three-dimensional light-sheet fluorescence microscopy as previously described(Ertürk et al., 2012). Tissue clearing was performed using the 3DISCO protocol, followed by imaging of intact brains with a light-sheet fluorescence microscope to visualize GFP-expressing GL261-gp100 tumor cells and mCherry-expressing Pmel- T cells. Three-dimensional image reconstruction and rendering were performed using Imaris software (version 9.3).

### Immunofluorescence staining

For immunofluorescence staining, formalin-fixed, paraffin-embedded tissue sections (3 µm) were deparaffinized in xylene (2 × 10 min) and rehydrated through a descending ethanol series (99% for 5 min, 96% for 2 min, and 70% for 2 min), followed by distilled water for 5 min. Heat-induced epitope retrieval was performed in a Tris-based buffer (pH 8.4; Discovery CC1, Roche) at 95 °C for 64 min, after which slides were allowed to cool in the same buffer for 30 min at room temperature. Sections were washed sequentially in 1× TBS (5 min) and in TBS containing 0.05% Tween-20 (TBST; 5 min), and non-specific binding was blocked with 10% normal goat serum (NGS; Gibco) in TBS for 30 min at room temperature.

Sections were then co-incubated with a rat anti-mouse CD8a antibody (clone 4SM15, 1:100; #14-0195-82, Invitrogen) and a rabbit anti-mouse CD31/PECAM-1 antibody (clone D8V9E, 1:68; #77699, Cell Signaling Technology), both diluted in 10% NGS/TBS, for 32 min at 37 °C. After three washes of 10 min each in TBST, bound primary antibodies were detected with goat anti-rat Alexa Fluor 546 (1:200; #A-11081, Invitrogen) and goat anti-rabbit Alexa Fluor 633 (1:200; #A-21070, Invitrogen) secondary antibodies diluted in 10% NGS/TBS and applied for 30 min at room temperature in the dark. Sections were washed three times for 5 min in TBST and rinsed for 1 min in MilliQ water. Nuclei were counterstained and sections were mounted using VECTASHIELD HardSet Antifade Mounting Medium with DAPI (#H-1500, Vector Laboratories) according to the manufacturer’s instructions, followed by coverslipping. Stained sections were scanned on an Axio Scan.Z1 slide scanner (Carl Zeiss) using a 20× objective.

For quantification of CD8⁺ T cell infiltration, whole-slide scans were imported into QuPath (v0.7.0-x64). Tumor regions were manually annotated, and cell detection was performed within these regions based on DAPI nuclear staining using the built-in positive cell detection algorithm. Detected cells were classified as CD8⁺ using a single-measurement object classifier based on maximum Alexa Fluor 546 (CD8^+^) intensity per cell, with the threshold adjusted individually for each slide. The percentage of CD8⁺ T cells was calculated as the number of AF546⁺ cells divided by the total number of DAPI⁺ cells within the annotated tumor area.

### Isolation of tumor and lung infiltrating immune cells

Mice were anesthetized and perfused with PBS. Brains of tumor-bearing mice were extracted, cerebellum and olfactory bulbs were removed, and the tumors were isolated from the right-hemisphere. The other hemispheres were used as a control. Whole lungs were isolated. The organs were further cut into pieces and digested with 50 μg/mL liberase (Sigma) at 37°C for 30 min and subsequently mashed through a 100 and 70 μm smart-strainer set (Miltenyi). Myelin and debris were removed using debris removal solution according to the manufacturer guidelines (Miltenyi).

### Fluorescence-activated cell sorting of tumor infiltrating murine T cells, myeloid cells and endothelial cells for single-cell RNA sequencing

Cell suspensions were blocked with anti-CD16/CD32 antibodies (eBioscience). eFluor 780 fixable viability dye (eBioscience) was used according to manufacturer’s protocol to exclude dead cells. Cells were sorted on a BDAria II through a100 μM nozzle and 4-way purity in TCM. The panel included Brilliant Violet (BV) 421 for CD3^+^ population, BV510 for CD45^+^ population, FITC for CD45.2^+^ population, PE for CD31^+^ population, APC for CD11b^+^ population, and PE-Cy7 CD45.1^+^ population. Unbiased mRNA profiling was then performed by using the chromium single-cell 5’ TCR/RNA sequencing kit with feature barcode technology (10x Genomics) as per manufacturer’s protocol. For cell multiplexing, samples were stained with TotalSeq-C hashtag antibodies. Hashtag oligonucleotide counts were used during downstream analysis for sample demultiplexing and doublet identification.

V(D)J repertoire sequencing was performed on the 10x Genomics Chromium platform. Brain and lung samples from four experimental groups (ACT, R-ACT, V-ACT, and V-R-ACT) were processed across multiple GEM wells. For surface protein detection, cells were stained using a custom TotalSeq-C mouse lyophilized panel (BioLegend). Briefly, lyophilized antibody panels were equilibrated to room temperature for 5 minutes and spun down at 10,000 × g for 30 seconds. Panels were reconstituted in 27.5 µL Cell Staining Buffer (BioLegend, Cat. No. 420201), vortexed for 10 seconds, and incubated at room temperature for 5 minutes. Following a second centrifugation step at 14,000 × g for 10 minutes at 4°C, 25 µL of the reconstituted cocktail was transferred to FcR-blocked cells (Mouse TruStain FcX™), with a final staining volume of 50 µL per 5 × 10⁵ cells.

GEX, CSP, and V(D)J libraries were prepared according to the 10x Genomics user guide for TotalSeq-C panels. Libraries were pooled and sequenced on a NovaSeq 6000 (Paired-End 150 bp, S4 flow cell, 600 Gbp/lane) targeting 8,000 million reads per pool, with a target of ≥85% Q30.

### Single-cell RNA sequencing analysis

Sequencing reads were processed with the Cell Ranger multi pipeline (v7.1.0), which jointly processed gene expression, cell hashing, and V(D)J sequencing data, and were aligned to the GRCm38 mouse reference genome. Downstream analysis was performed in R (v4.5.1) using Seurat (v5.4.0); visualizations were generated with SCpubr (v3.0.1) and ggplot2 (v4.0.3).

Cells with >15% mitochondrial reads, or feature/UMI counts below 30% (GL261-OVA) or 20% (GL261-gp100) of the sample median, were excluded; upper thresholds were set at the mean + 3 s.d. Data were log-normalized, and 2,000 variable features were used to scale the data while regressing out mitochondrial and ribosomal content. PCA was performed on variable features, and the first 50 principal components were kept as chosen based on the elbow plot of explained variance. These principal components were used for clustering (shared nearest-neighbor graph, resolution = 1.0), followed by UMAP visualization. T cell and endothelial cell subsets were re-processed independently using the same workflow, except for the GL261-OVA T cell subset, which was clustered at a higher resolution (1.9) for a more granular annotation.

Clusters were annotated using literature-supported marker genes derived from differential expression testing between clusters. Endothelial subclusters were further assigned identities by mapping to reference signatures from Xie et al.(Xie et al., 2021), based on the highest enrichment score per cluster.

Genes were tested for differential expression between therapy conditions within each cell type, restricted to genes detected in ≥30% of cells in at least one group (min.pct = 0.3); significance was defined as adjusted P < 0.05. Enrichment of biological process terms was performed with clusterProfiler (v4.4.4), mapping genes to Ensembl IDs via org.Mm.eg.db (3.22.0), and visualized with enrichplot (v1.16.2).

Predefined (Table X) and differential-expression–derived (Table Y) gene signatures were scored per cell using Seurat’s module-scoring approach. Mean enrichment across conditions was visualized as a heatmap (SCpubr, “Seurat” flavor), ranked by descending mean score. Per-replicate averaged scores were compared between treatment groups using boxplots and two-sided t-tests (ggpubr, v0.6.2).

Clonotypes from Cell Ranger VDJ output were merged across samples and integrated into the T cell Seurat object with scRepertoire (2.6.2), assigning each cell a clonal expansion category (Rare ≤1×10⁻⁴; Small ≤0.001; Medium ≤0.01; Large ≤0.1; Hyperexpanded ≤1). Clonal diversity per biological replicate was estimated using Shannon entropy (amino-acid level, 100 bootstraps) and compared across conditions.

Ligand–receptor interactions were inferred using LIANA (0.1.14) with the OmniPath consensus resource, independently per therapy condition; interactions with an aggregate rank ≤0.01 were retained. Cross-condition comparisons were restricted to interactions detected in both conditions (inner join) with a minimum magnitude of 0.3 in at least one condition, with differential activity quantified as the difference in sca.LRscore. Communication was additionally assessed using CellChat (v1.6.1), with selected interactions visualized as chord diagrams.

### Analysis of human spatial transcriptomic data

Xenium spatial transcriptomics data from (Seferbekova et al., 2026) were analyzed from preprocessed AnnData object as follows: Cells annotated as ’uninformative cells’ were excluded. UMAP coordinates were recomputed from the provided scANVI representation (15 nearest neighbors). Provided cell type annotations were validated using curated marker gene dot plots. Endothelial cells were re-clustered independently using Leiden clustering (resolution = 0.8) on the scANVI embedding. Clusters were annotated into subpopulations (Co1, Co2, Co3, Pe1, Pe2) by scoring published endothelial reference signatures (Xie et al., 2021) using scanpys ‘score_genes’ function. T cells were similarly re-clustered (Leiden, resolution = 0.5) and annotated as CD4^+^ or CD8^+^ T cells based on canonical markers (CD4, IL7R, CD8A, CD8B).

Gene set scores were computed with ‘sc.tl.score_genes’ for positive regulation of type I interferon production (GO:0032481), interferon-mediated signaling (GO:0140888), and aggregate HLA gene expression. Differentially expressed genes in endothelial cells between RI and non-RI patients were identified by Wilcoxon rank-sum test on scVI-normalized values (adjusted p < 0.01, |log fold change| > 0.2). Cross-species comparison with externally provided mouse endothelial radiation DEGs was performed by converting mouse gene symbols to human homologs via the MyGeneInfo API (HomoloGene-based).

Spatial co-occurrence of cell type pairs was computed using Squidpy (‘sq.gr.co_occurrence’) per condition over 0–300 µm (10 µm steps). Neighborhood enrichment z-scores were calculated using Delaunay triangulation-based neighborhood graphs (‘sq.gr.nhood_enrichment’) per condition, and the difference between conditions was visualized as a heatmap. For each patient, endothelial cell positions were binned onto a 2D grid (20 µm/pixel). The density grid was smoothed with a Gaussian filter (σ = 1.5 pixels) and thresholded (threshold = 0.4) to generate a binary vessel mask. Connected components were labeled as individual vessels using ‘scipy.ndimage.label’. The Euclidean distance transform was applied to the inverted mask to compute each cell’s distance to the nearest vessel boundary. All distances were converted to micrometers using the Xenium scale factor (0.2125 µm/pixel). Each cell was assigned a binary in-vessel flag, a vessel identifier, and a continuous distance value. Cell type composition per vessel was quantified as the proportion of each cell type among all cells within a vessel, and compared between RI and non-RI conditions using the Mann-Whitney U test (two-sided). Cells were additionally binned by distance from vessels (0–20, 20–50, 50–100, 100–200, >200 µm) and cell type proportions within each bin were compared between conditions using the Mann-Whitney U test.

### Statistical analysis

Data are presented as mean ± SEM unless otherwise indicated. Group sizes (n) and statistical tests are provided in the figure legends. Statistical analyses were performed using GraphPad Prism and R (v4.4.1). Survival was analyzed using the Kaplan–Meier method and compared using the log-rank test. Differences between groups were assessed using unpaired or paired *t*-tests or one-way ANOVA with Tukey’s post hoc test, as appropriate. Adjusted *P* values were corrected using the Benjamini–Hochberg method where applicable.

## Acknowledgements

We thank Manuel Fischer for expert assistance with magnetic resonance imaging (MRI), Julia Bode for excellent technical support, and Björn Tews for support with tissue clearing and light-sheet fluorescence microscopy. We are grateful to all members of our laboratories for helpful discussions and technical assistance throughout the course of this study.

## Declaration of generative AI and AI-assisted technologies in the manuscript preparation process

During the preparation of this work the authors used ChatGPT (OpenAI) and Claude (Anthropic) in order to assist with computational data analysis and text drafting. After using this tool/service, the authors reviewed and edited the content as needed and take full responsibility for the content of the published article.

## Author Contributions

CTD, NG, NE, MP, LB, and KS conceived and designed the study. NE, MOB, DAA, TB, KJ, JKS, RS and KhS performed experiments and acquired data. NE established, and executed the interventional pipelines for single-cell analysis in murine models. AS, MG and FS provided and interpreted human data. CTD, NG, NE and KS analyzed data. CTD, NG, JR, AM and ZS performed computational and bioinformatic analyses. CTD, NG and KS interpreted the results and developed the conceptual framework of the study. MP and KS supervised the research. MP, LB and KS acquired funding. CTD, NG, NE, and KS wrote the manuscript with input from all authors. All authors reviewed, edited, and approved the final manuscript.

